# C-Jun N-terminal Kinase Promotes Stress Granule Assembly and Neurodegeneration in C9orf72-mediated ALS and FTD

**DOI:** 10.1101/2022.04.28.489917

**Authors:** TG Sahana, Katherine Johnson Chase, Feilin Liu, Thomas E. Lloyd, Wilfried Rossoll, Ke Zhang

**Affiliations:** Department of Neuroscience, Mayo Clinic, Jacksonville, FL 32224, USA; Department of Neurology, Johns Hopkins School of Medicine, MD 21205, USA; Neuroscience Graduate Program, Mayo Clinic Graduate School of Biomedical Sciences, Jacksonville, FL 32224, USA

## Abstract

Stress granules (SGs), RNA/protein condensates assembled in cells under stress, are believed to play a critical role in the pathogenesis of amyotrophic lateral sclerosis (ALS) and frontotemporal dementia (FTD). However, how SG assembly is regulated and related to pathomechanism is incompletely understood. Here, we show that ER stress activates JNK via IRE1 in fly and cellular models of C9orf72-mediated ALS/FTD (c9ALS/FTD), the most common genetic form of ALS/FTD. Furthermore, activated JNK promotes SG assembly induced by poly(GR) and poly(PR), two toxic proteins implicated in c9ALS/FTD, by promoting the transcription of G3BP1, a key SG protein. Consistent with these findings, JNK or IRE1 inhibition reduced SG formation, G3BP1 mRNA and protein levels, and neurotoxicity in cells overexpressing poly(GR) and poly(PR) or neurons derived from c9ALS/FTD patient induced pluripotent stem cells (iPSCs). Our findings connect ER stress, JNK, and SG assembly in a unified pathway contributing to c9ALS/FTD neurodegeneration.

## Introduction

A GGGGCC (G_4_C_2_) hexanucleotide repeat expansion in chromosome 9, open reading frame 72 (*C9ORF72*) is the most common genetic cause of amyotrophic lateral sclerosis (ALS) and frontotemporal dementia (FTD) (DeJesus-Hernandez et al., 2011; Renton et al., 2011). This repeat expansion can cause cytotoxicity via multiple mechanisms, one of which suggests that it undergoes repeat-associated, non-ATG translation to produce five different species of dipeptide repeat proteins (DPRs), namely poly(glycine-arginine, GR), poly(glycine-alanine, GA), poly(glycine-proline, GP), poly(proline-alanine, PA), and poly(proline-arginine, PR)(Ash et al., 2013; Donnelly et al., 2013; Gendron et al., 2013; Ling, Polymenidou, & Cleveland, 2013; Mori, Arzberger, et al., 2013; Mori, Weng, et al., 2013; Zu et al., 2013). Among these DPRs, the arginine-rich DPRs (R-DPRs) i.e., poly(GR), and poly(PR), are especially toxic (Kwon et al., 2014; K. H. Lee et al., 2016; Lin et al., 2016; Mizielinska et al., 2014; Sakae et al., 2018; K. Zhang et al., 2018; Y. J. Zhang et al., 2018; Zhang et al., 2019).

Stress granules (SGs) are cytoplasmic RNA/protein condensates assembled in cells under stress (Protter & Parker, 2016). Upon stress, polysomes disassemble, and mRNAs are embedded by a variety of RNA-binding proteins, whose condensation mediates SG assembly (Guillen-Boixet et al., 2020; Sanders et al., 2020; P. Yang et al., 2020). Under normal conditions, SGs are dynamic and disassemble when stress is removed (Lin, Protter, Rosen, & Parker, 2015; Protter & Parker, 2016). However, aberrant SG formation can trigger aggregation of SG proteins, such as TDP-43 and FUS (A.N. Coyne et al., 2015; Daigle et al., 2013). Since the aggregation of these proteins is a pathological hallmark of ALS and FTD, including c9ALS/FTD, SGs are believed to play a critical role in ALS/FTD pathogenesis (Anderson & Kedersha, 2008; Kedersha, Tisdale, Hickman, & Anderson, 2008; Li, King, Shorter, & Gitler, 2013). Consistent with this notion, R-DPRs interact with many SG proteins, and their overexpression causes the formation of aberrant, poorly dynamic SGs in cells without additional stress (Boeynaems et al., 2017; K. H. Lee et al., 2016; K. Zhang et al., 2018). In addition, chemically synthesized R-DPRs can undergo liquid-liquid phase separation (LLPS), recruit SG proteins, and cause SG protein precipitation in cellular lysates (Boeynaems et al., 2017). Also, poly(GR) can localize to SGs, promote the aggregation of recombinant TDP-43 *in vitro*, and co-aggregate with TDP-43 and the SG protein eIF3η in c9ALS/FTD patient postmortem tissue (Cook et al., 2020). In agreement with these data, we previously found that inhibiting SG assembly by genetic or pharmacological approaches suppresses R-DPR-induced cytotoxicity or neurodegeneration in cellular or animal models (K. Zhang et al., 2018). Together, these findings suggest that R-DPRs cause neurodegeneration by promoting aberrant SG formation. However, how this process is regulated is unclear.

In a *Drosophila* RNAi screen, we previously identified that loss of *bsk*, the fly homolog of c-Jun N-terminal kinase (JNK), suppresses neurodegeneration in a fly model of c9ALS/FTD (K. Zhang et al., 2015). Here, we show that JNK is activated in fly and cellular models of c9ALS/FTD via ER stress response protein IRE1 and activated JNK promotes R-DPR-induced SG formation by promoting the transcription of G3BP1, a key protein involved in SG assembly (Deniz, 2020; Guillen-Boixet et al., 2020; P. Yang et al., 2020). Inhibiting ER stress responses or JNK activity suppresses R-DPR-induced SG formation, G3BP1 mRNA and protein levels, and cytotoxicity in cells expressing R-DPRs or c9ALS/FTD patient iPSC-derived neurons (iPSNs). Our findings identified a molecular mechanism by which the ER stress/IRE1/JNK axis promotes long-term-stress-induced SG formation and suggested a unified, druggable pathway contributing to c9ALS/FTD pathogenesis.

## Results

### Loss of *bsk/JNK* suppresses neurodegeneration caused by G_4_C_2_ repeats in *Drosophila*

Expression of 30 G_4_C_2_ repeats [(G_4_C_2_)_30_] in fly eyes using GMR-GAL4 causes neurodegeneration, as indicated by defects in the external eye morphology that worsen with age (Xu et al., 2013; K. Zhang et al., 2015). Using this fly model, our previously published RNAi screen identified *bsk* RNAi to potently suppress (G_4_C_2_)_30_-mediated eye degeneration (K. Zhang et al., 2015), which we verified (**Fig. 1a**). Furthermore, we show that (G_4_C_2_)_30_ expression in motor neurons using vGlut-GAL4 causes paralysis in pharate flies, as indicated by their inability to eclose (Cunningham et al., 2020). Here, we show that *bsk* RNAi suppresses this phenotype (**Fig. 1b**). Thus, loss of *bsk* suppresses (G_4_C_2_)_30_-mediated toxicity in fly eyes and motor neurons.

**Figure 1:**
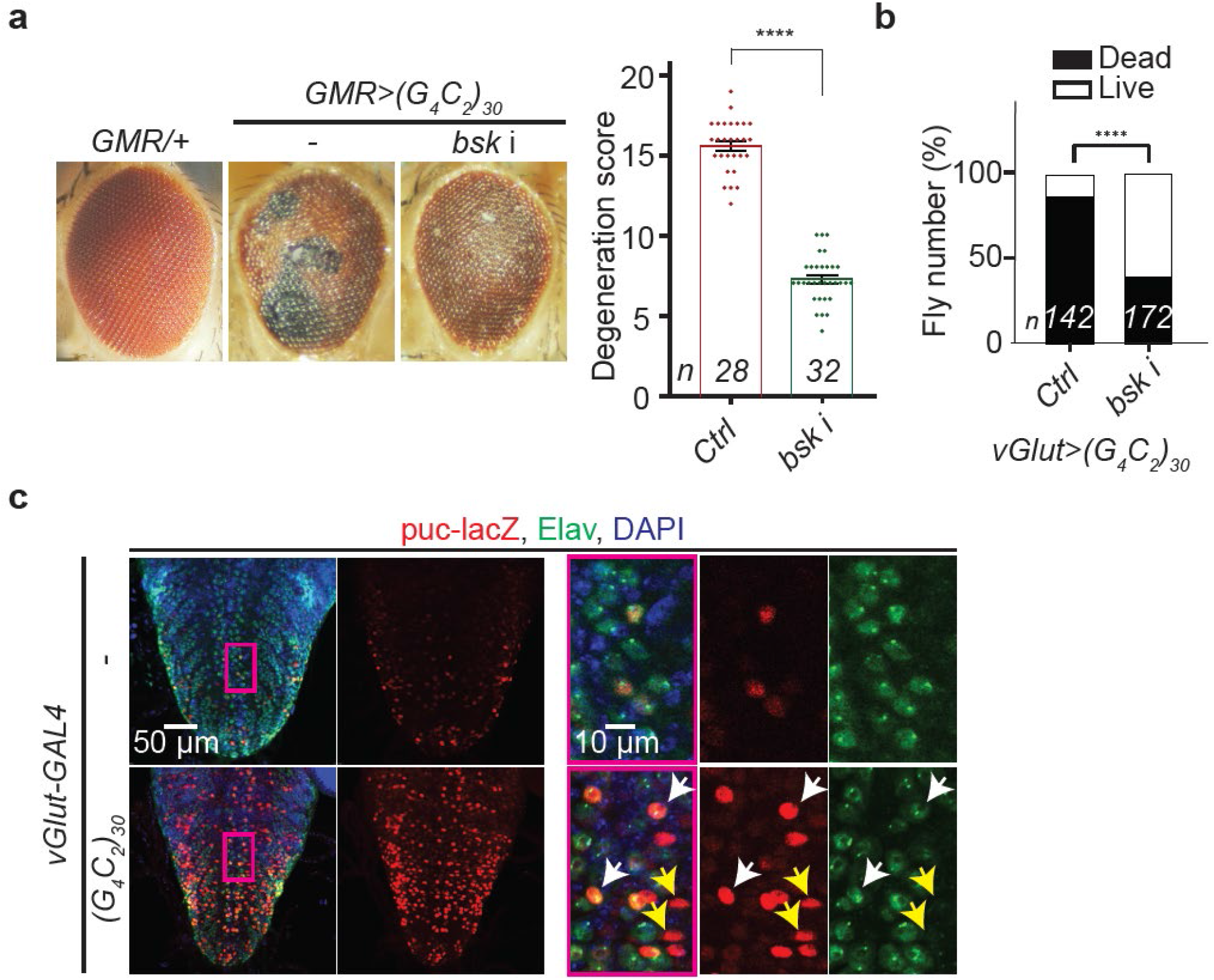
JNK/bsk is activated in a fly model of c9ALS/FTD. (a) Fly eyes expressing (G_4_C_2_)_30_ using GMR-GAL4, without or with *bsk* RNAi (*bsk* i). Scored using a previously published method (G.P. Ritson et al., 2010). Mean ± s.e.m. Student’s t-test; ****: *p*<0.0001 (b) Percent of eclosed adult flies expressing (G_4_C_2_)_30_ in motor neurons using vGlut-GAL4, without (control, Ctrl) or with *bsk* i. χ2-test; ****: *p*<0.0001 (c) Fly ventral nerve cord motor neurons expressing puc-lacZ without or with co-expressing (G_4_C_2_)_30_, stained with lacZ (red), Elav (a neuronal marker, green), and DAPI (blue).

Next, we tested whether Bsk is hyperactive in neurons expressing G_4_C_2_ repeats. Bsk/JNK belongs to the mitogen-activated protein kinase (MAPK) family, which is activated by JNK kinases (Kim & Choi, 2010; Sahana & Zhang, 2021). Upon activation, Bsk/JNK activates the transcription of downstream genes, including its inhibitor, JNK phosphatase (fly homolog: *puckered*, or *puc*). Thus, the level of Puc/JNK phosphate can be used to indicate Bsk/JNK activity. Indeed, a LacZ reporter under the control of the *puc* promoter (puc-LacZ) is widely used as a reporter of Bsk/JNK activity in flies (Martin-Blanco et al., 1998; Ring & Arias, 1993). Using this reporter, we show that the LacZ level is strongly upregulated in motor neurons expressing (G_4_C_2_)_30_, compared to the control (**Fig. 1c**), suggesting that expression of G_4_C_2_ repeats causes JNK activation.

### Expression of G_4_C_2_ repeats causes ER stress in *Drosophila*

Previous studies showed that JNK can be activated by ER stress. Upon ER stress, inositol requiring enzyme 1 (IRE1), a protein with both kinase and endonuclease activities, is activated, which recruits the tumor necrosis factor receptor-associated factor 2 (TRAF2). The IRE1/TRAF2 complex phosphorylates and activates apoptosis signal-regulating kinase (ASK-1), a MAP kinase kinase kinase (MAP3K), which activates its downstream target JNK (Urano et al., 2000). Furthermore, previous studies have implicated ER stress in c9ALS/FTD iPSNs (Dafinca et al., 2016; Haeusler et al., 2014). Here, we show that RNAi against IRE1, TRAF2, or ASK-1 suppresses eye degeneration in flies expressing (G_4_C_2_)_30_, suggesting that these proteins contribute to G_4_C_2_-repeat-mediated neurotoxicity (**Fig. 2a**).

**Figure 2:**
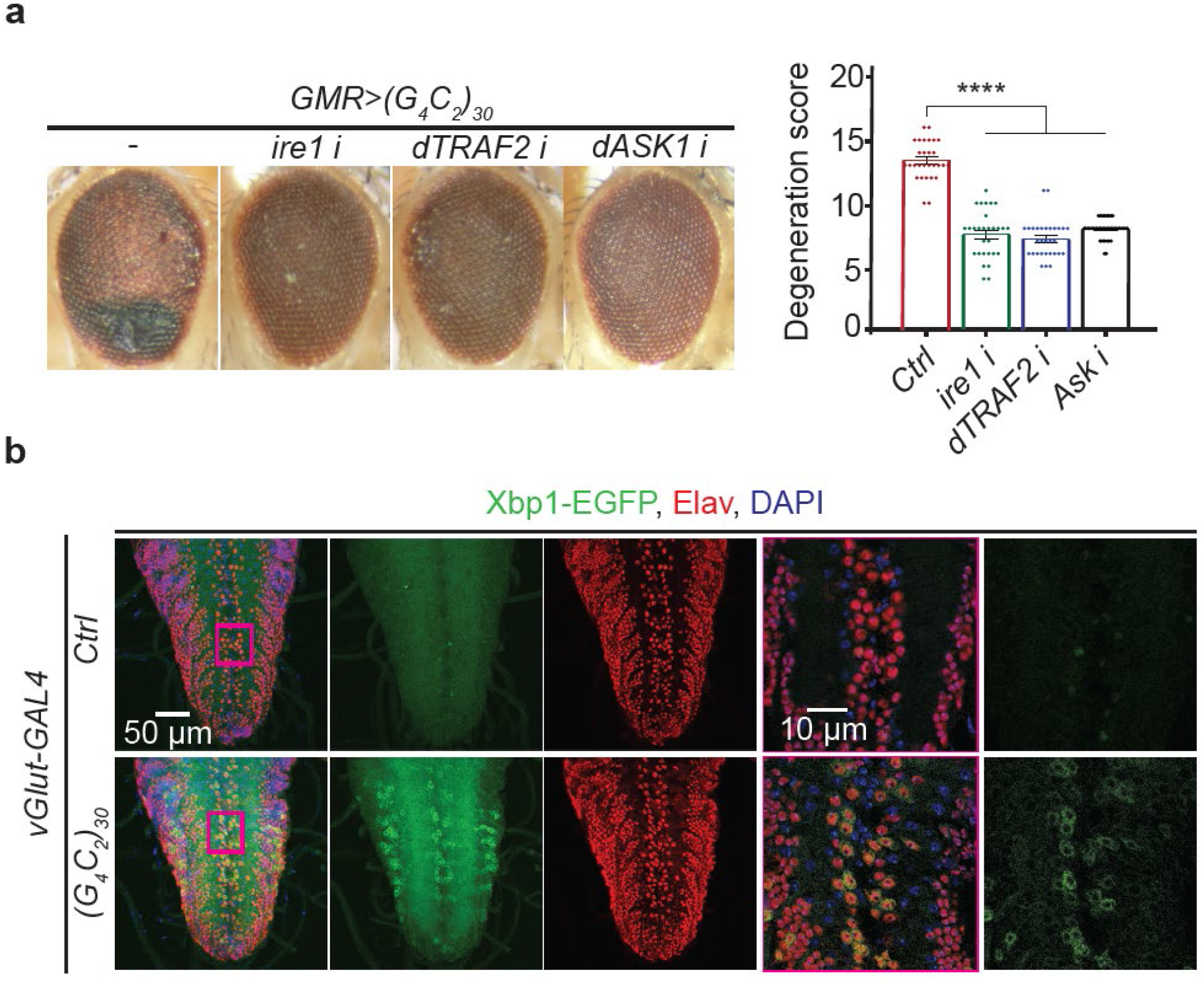
ER stress contributes to neurodegeneration in a fly model of c9ALS/FTD. (a) Fly eyes expressing (G_4_C_2_)_30_ without or with RNAi against ire1, dASK1 and dTRAF2. Mean ± s.e.m. One-way ANOVA; ****: *p*<0.0001. (b) Fly VNC motor neurons expressing Xbp1-GFP without or with (G_4_C_2_)_30_, stained with Elav (a neuronal marker, red) and DAPI (blue).

Next, we investigated the IRE1 activity in fly neurons. Upon IRE1 activation, its endonuclease activity causes the alternative splicing of X-box-binding protein 1 (XBP1) mRNA. In fly studies, a widely used IRE1 reporter is an XBP1-GFP system, in which GFP is expressed only when XBP1 is alternatively spliced due to IRE1 activation (Sone, Zeng, Larese, & Ryoo, 2013). Using this system, we show that GFP is strongly upregulated in motor neurons expressing (G_4_C_2_)_30_, compared to the control (**Fig. 2b**), suggesting that expression of G_4_C_2_ repeats causes IRE1 activation. Together, our data suggest that G_4_C_2_ repeats cause ER stress, further leading to toxicity in *Drosophila*.

### R-DPRs cause JNK activation and ER stress in U-2 OS cells

Among five DPR species, the R-DPRs, i.e., poly(GR) and poly(PR), are highly toxic and cause eye degeneration and cytotoxicity in *Drosophila* and cultured cells (Mizielinska et al., 2014). Thus, we investigated their roles in JNK activation and ER stress. First, we show that *bsk* RNAi suppresses eye degeneration caused by 36 repeats of poly(GR) or poly(PR) (**Supplementary Fig. 1**), suggesting that JNK contributes to poly(GR) and poly(PR)-mediated toxicity. Next, we switched to U-2 osteosarcoma (OS) cells, which are widely used to study cellular stress responses and R-DPR toxicity for their human relevance and ease to dissect cellular mechanisms (Boeynaems et al., 2017; Kwon et al., 2014; Ohn, Kedersha, Hickman, Tisdale, & Anderson, 2008; P. Yang et al., 2020).

Using an MTT cell survival assay, we show that transiently expressing mCherry-tagged, 100 repeats of poly(GR) or poly(PR) [(GR)_100_- or (PR)_100_-mCherry] for 48 hours impairs U-2 OS cell survival, compared to the mCherry control, which is partially suppressed by a 24-hour co-treatment of a pan-JNK inhibitor, SP600125 (**Fig. 3a**). These data suggest that inhibiting JNK activity suppresses R-DPR-mediated cytotoxicity, consistent with our fly data (**Supplementary Fig. 1**). Furthermore, both our immunofluorescent staining and Western blots show that (GR)_100_- and (PR)_100_-mCherry increase the levels of phosphorylated JNK (pJNK), the activated JNK form, compared to the mCherry control (**Fig. 3b and c**), suggesting that R-DPRs activate JNK in U-2 OS cells.

**Figure 3:**
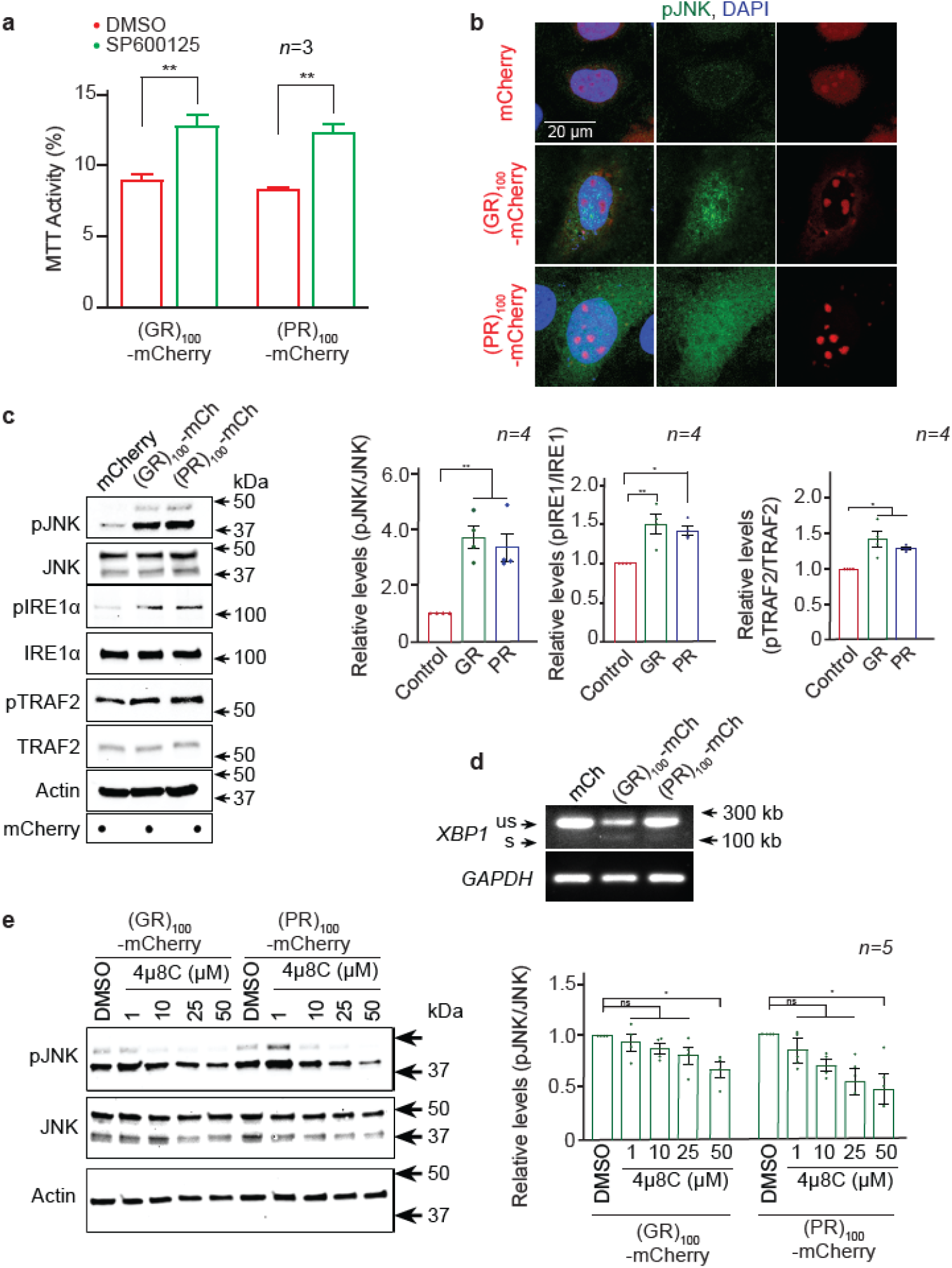
ER stress and JNK is activated in U-2 OS cells expressing R-DPRs. (a) MTT assays of U-2 OS cells expressing (GR/PR)_100_-mCherry, treated with DMSO or 50 µM of SP600125 for 24 h. MTT activity of U-2 OS cells expressing mCherry is taken as 100%. Mean ± s.e.m. Student’s t-tests; **, *p*<0.01. (b) U-2 OS cells expressing mCherry or (GR/PR)_100_-mCherry (red) stained with pJNK (green) and DAPI (blue). (c) Western or dot (for mCherry only) blots for lysates from U-2 OS cell expressing mCherry or (GR/PR)_100_-mCherry. Mean ± s.e.m. One-way ANOVA; **, *p*<0.01; *, *p*<0.05. (d) DNA gels showing spliced variants of XBP1 (us: unspliced and s: spliced) from cDNA of U-2 OS cell expressing mCherry or (GR/PR)_100_-mCherry. (e) Western blots for lysates from U-2 OS cell expressing mCherry or (GR/PR)_100_-mCherry co-treated with DMSO or 4µ8C for 6 h. Mean ± s.e.m. One-way ANOVA; *, *p*<0.05, ns, not significant.

Next, we investigated whether (GR)_100_- or (PR)_100_-mCherry induces ER stress. As shown in **Fig. 3c and d**, transient expression of (GR)_100_- or (PR)_100_-mCherry causes upregulation of phosphorylated IRE1 (pIRE1) and TRAF2 (pTRAF2), i.e., activated forms of IRE1 and TRAF2, as well as alternative splicing of XBP1 mRNA in U-2 OS cells, compared to the mCherry control, suggesting that R-DPRs activates IRE1 in these cells. Of note, 4µ8C, an inhibitor of both IRE1 kinase and endonuclease activities (Cross et al., 2012), dose-dependently suppresses pJNK levels in U-2 OS cells expressing (GR)_100_- or (PR)_100_-mCherry (**Fig. 3e**), suggesting that R-DPRs activate JNK via IRE1.

### Inhibiting JNK or IRE1 activity suppresses R-DPR-induced SG formation in U-2 OS cells

SGs play a critical role in R-DPR-mediated cytotoxicity. Indeed, R-DPRs can induce poorly dynamic SGs in cultured cells without additional stress (Boeynaems et al., 2017; K. H. Lee et al., 2016). When expressed in U-2 OS cells for 48 hours, (GR)_100_- or (PR)_100_-mCherry respectively causes ∼70% or ∼30% cells to exhibit SGs, as indicated by immunofluorescent staining of G3BP1 and TIA1, two SG markers (**Fig. 4** and **Supplementary Fig. 2**). Furthermore, treating the cells with 50 µM of SP600125 for 24 hours or 4µ8C for six hours significantly decreases the percent of cells exhibiting SGs, but not R-DPR protein levels (**Fig. 4** and **Supplementary Fig. 2**), suggesting that IRE1/JNK promotes R-DPR-induced SG assembly.

**Figure 4:**
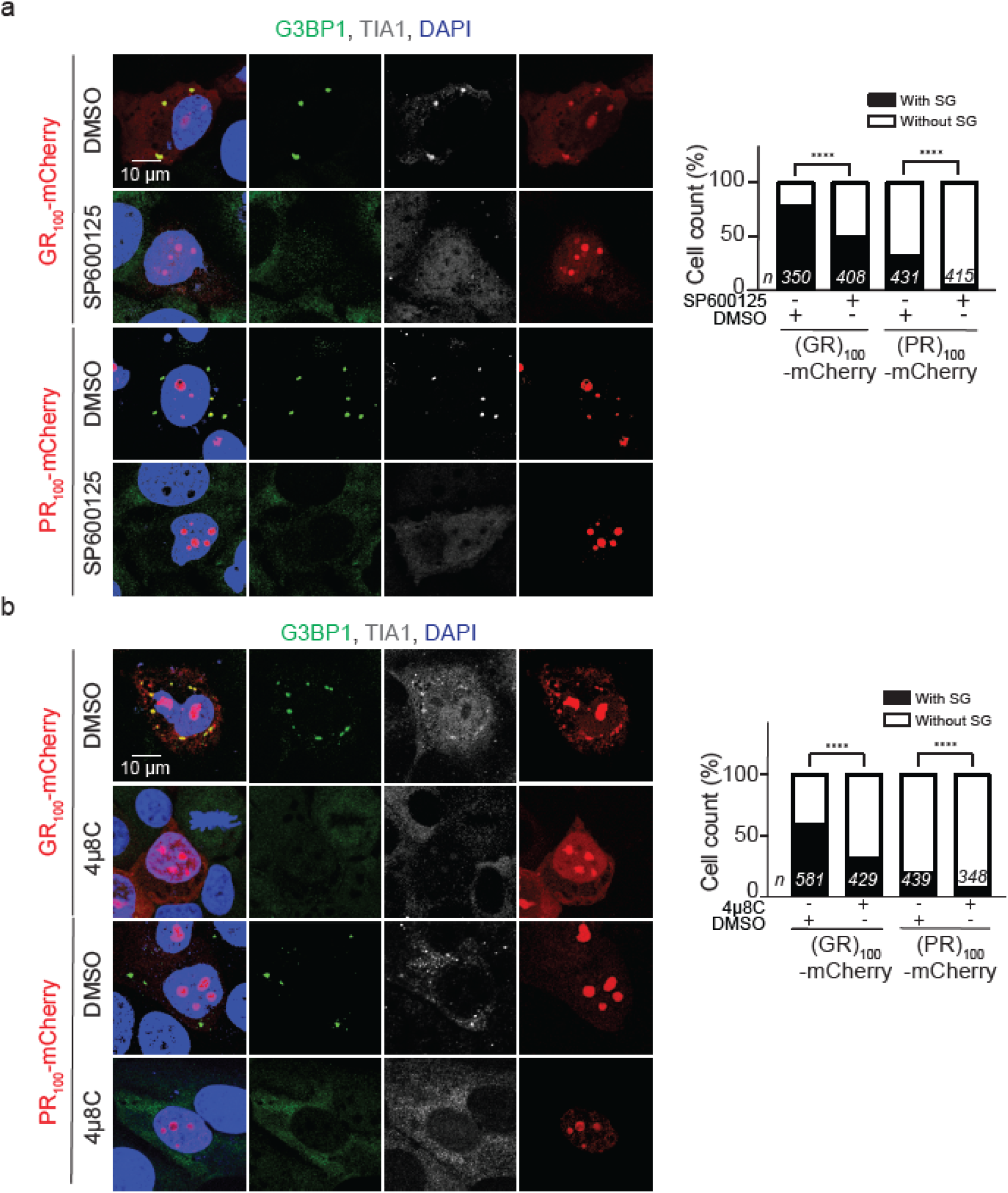
ER stress and JNK promote R-DPR-induced SG assembly in U-2 OS cells. U-2 OS cells expressing (GR/ PR)_100_-mCherry (red) treated with DMSO, 50 µM SP600125, or 50 µM 4µ8C and stained with G3BP1 (green), TIA1 (white), and DAPI (blue). Quantification showing percent of cells with or without SGs. χ2-test; ****: *p*<0.0001.

G3BP1 plays a critical role in SG assembly, as its knockdown strongly reduces SG formation caused by a variety of stressors, whereas its overexpression induces SG formation without additional stress (Kedersha et al., 2016). In addition, we and others found that double-knockout of G3BP1 and its homolog, G3BP2, completely abolishes R-DPR-induced SGs (Boeynaems et al., 2017; K. Zhang et al., 2018). Here, we show that a 24- or six-hour treatment of SP600125 or 4µ8C, respectively, strongly decreases G3BP1 levels in cells expressing (GR)_100_- or (PR)_100_-mCherry (**Fig. 5a and b**), suggesting that JNK inhibition downregulates G3BP1. In addition, we found that these inhibitors decrease G3BP2, but not TIA1, levels (**Fig. 5a**), suggesting that JNK regulates some SG proteins.

**Figure 5:**
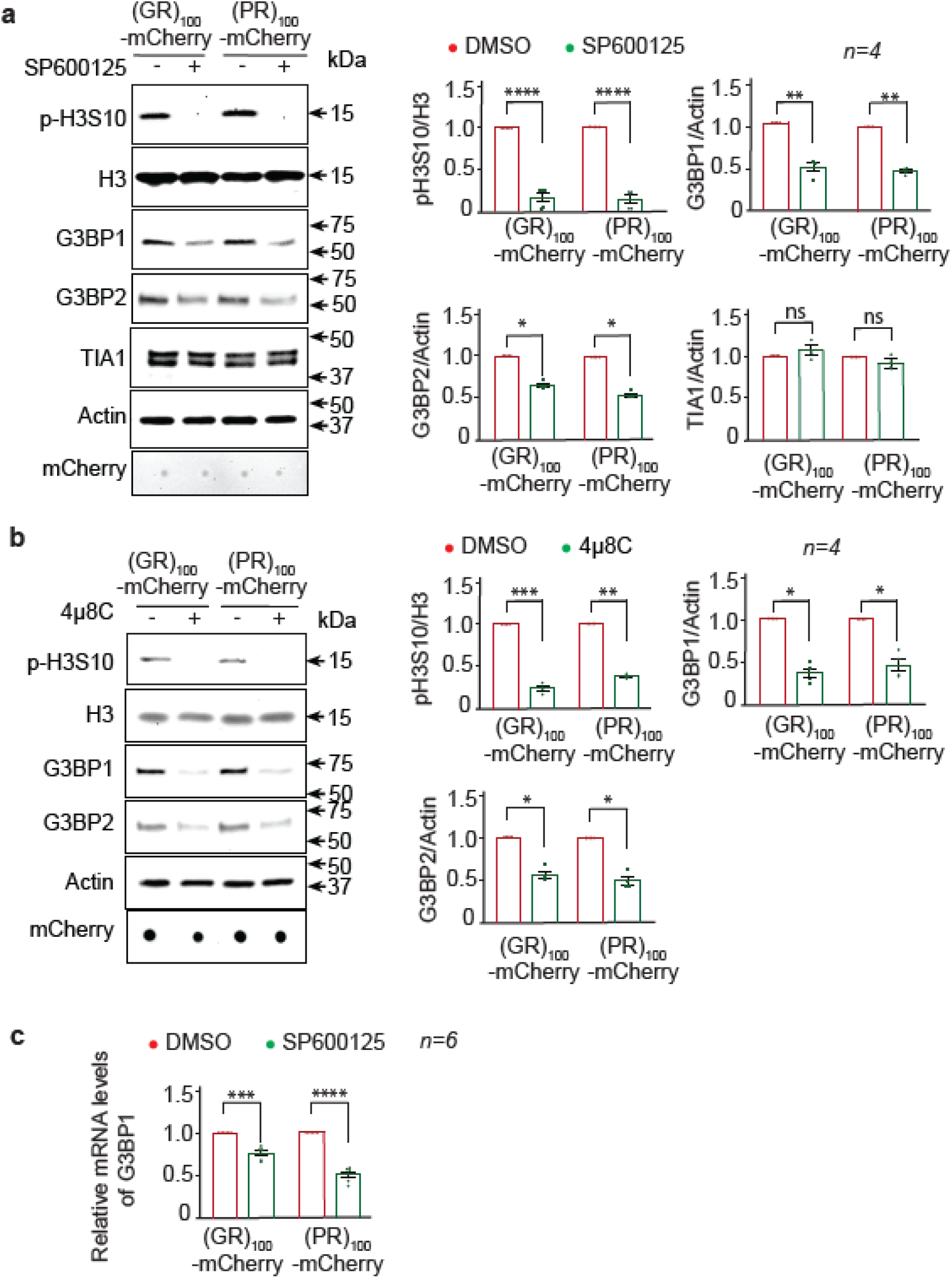
JNK promotes G3BP1 expression and H3S10 phosphorylation in U-2 OS cell expressing R-DPRs. Western or dot (for mCherry only) blots of lysates from U-2 OS cells expressing (GR/PR)_100_-mCherry, treated with DMSO or (a) 50 µM SP600125 and (b) 50 µM 4µ8C. (c) Relative levels of G3BP1 mRNA as compared to GAPDH from U-2 OS cells expressing (GR/PR)_100_-mCherry, treated with DMSO or 50 µM SP600125. Mean ± s.e.m. Student’s t-tests; ****, *p*<0.0001; ***, *p*<0.001; **, *p*<0.01; *, *p*<0.05; ns, not significant.

A U-2 OS cell line stably expressing GFP-tagged G3BP1 (G3BP1-GFP) under the control of a lentiviral promoter is widely used to study SG biology (Figley, Bieri, Kolaitis, Taylor, & Gitler, 2014). We found that SP600125 suppresses the level of endogenous G3BP1, but not G3BP1-GFP, in these cells when transfected with (GR)_100_- or (PR)_100_-mCherry (**Supplementary Fig. 3**), possibly because the regulation of JNK on G3BP1 relies on the genomic promoter of *G3BP1*. If this is the case, JNK inhibition likely suppresses *G3BP1* transcription. Indeed, as shown in **Fig. 5c**, SP600125 significantly decreases *G3BP1* mRNA levels in U-2 OS cells expressing (GR)_100_- or (PR)_100_-mCherry, suggesting that JNK regulates G3BP1 at the transcriptional level in these cells.

Widely used in SG studies, sodium arsenite induces ER stress and SG assembly within an hour (Anderson & Kedersha, 2008; Cheng et al., 2018; Wheeler, Matheny, Jain, Abrisch, & Parker, 2016; P. Yang et al., 2020). To test whether JNK also plays a role in arsenite-induced SG formation, we co-treated U-2 OS cells with 0.5 mM sodium arsenite and 50 µM of SP600125 for an hour. As shown in **Supplementary Fig. 4**, SP600125 suppresses the pJNK level, but not the percentage of SG-positive cells, suggesting that JNK does not contribute to SG assembly caused by one-hour arsenite stress. Consistent with these data, cellular G3BP1 levels are unaltered in these cells, suggesting that one-hour JNK inhibition is insufficient to affect G3BP1 protein levels or SG assembly.

A prior study showed that in mouse differentiating neurons, activated JNKs in the nucleus are enriched in the promoter regions of certain genes, including G3BP1, where they phosphorylate certain chromatin components i.e., histone 3 protein at Serine10 position (H3S10) (Tiwari et al., 2011). As H3S10 phosphorylation (pH3S10) causes the chromatin to adopt an “open” chromatin structure, which activates transcription (Allis & Jenuwein, 2016; Rossetto, Avvakumov, & Cote, 2012; Stricker, Koferle, & Beck, 2017), one possible mechanism by which JNK regulates *G3BP1* transcription is via pH3S10. Consistent with this notion, a 24- or six-hour treatment of SP600125 or 4µ8C, respectively, strongly suppresses pH3S10 levels in U-2 OS cells expressing (GR)_100_- or (PR)_100_-mCherry **(Fig. 5a and b)**.

### Inhibiting IRE1/JNK activity suppresses neurotoxicity in c9ALS/FTD patient-derived iPSNs

To validate our findings in a patient-relevant model, we used iPSC-derived neurons (iPSNs) derived from c9ALS/FTD patients. While these iPSNs rarely exhibit SGs under non-stressed conditions, we previously showed that they are constitutively under a low level of stress, as indicated by a mild increase in the phospho-eIF2α. In addition, SG inhibitors GSK2606414 and ISRIB, which suppress R-DPR-induced SG formation, suppress subcellular defects in these iPSNs, suggesting that SG assembly contributes to the iPSN toxicity (K. Zhang et al., 2018).

Compared to control iPSNs, c9ALS/FTD iPSNs do not exhibit a strong phenotype or reduced survival under non-stressed conditions but are more sensitive to a variety of stressors, including the ER stressor tunicamycin (Donnelly et al., 2013; Haeusler et al., 2014; Shi et al., 2018). We show that a 24-hour treatment of five µM tunicamycin (TM) causes cell death, as indicated by propidium iodide (PI) staining, in four different c9ALS/FTD iPSN lines, which is suppressed by co-treatments of either SP600125 or 4µ8C (**Fig. 6a and Supplementary Fig. 5**). Consistent with these data, we show that the G3BP1 level in the c9ALS/FTD iPSNs increases upon tunicamycin treatment, which is suppressed by SP600125 and 4µ8C **(Fig. 6b**). Together, these data suggest that inhibiting IRE1/JNK activity suppresses neurotoxicity and the G3BP1 level in c9ALS/FTD iPSNs.

**Figure 6:**
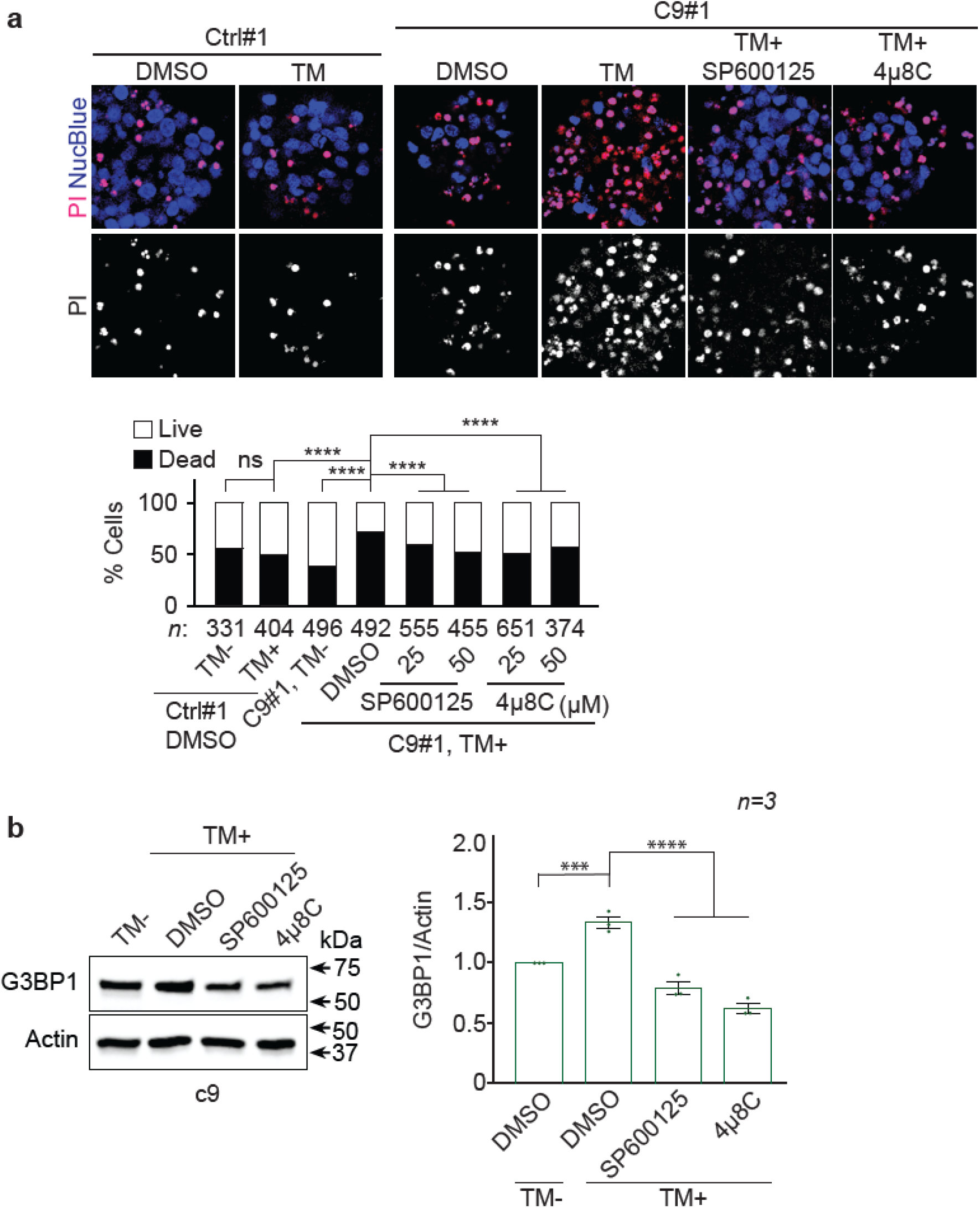
Inhibition of JNK or ER stress suppresses toxicity in c9ALS/FTD iPSNs. (a) Control (Ctrl) or c9ALS/FTD (c9) Line #1 iPSNs treated with 5 µM tunicamycin (TM) together with DMSO, SP600125, or 4µ8C and stained with propidine iodide (PI, dead cells) and NucBlue (all cells). Quantification shows the percent of live and dead cells. χ2-test. Mean ± s.e.m. One-way ANOVA with Dunnett’s test; ****: p<0.0001. (b) Western blot of c9 lysates treated with 5 µM tunicamycin (TM) together with DMSO, JNK inhibitor SP600125, or IRE1 inhibitor 4µ8C. Mean ± s.e.m. Student’s t-tests; ****, p<0.0001; ***, p<0.001.

### Loss of G3BP/Rin suppresses neurodegeneration in c9ALS/FTD fly models

Previously, we showed that G3BP1/2 double KO abolishes R-DPR-induced cellular defects in U-2 OS cells and SG inhibitors GSK2606414 and ISRIB suppress (G_4_C_2_)_30_-mediated eye degeneration in flies (K. Zhang et al., 2018), suggesting that inhibiting SG formation suppresses c9ALS/FTD-related cytotoxicity or neurodegeneration. Given the importance of G3BP1 and 2 in SG assembly, we postulate that loss of G3BP also suppresses neurodegeneration. In flies, *rin* is the only homolog of mammalian *G3BP1* and *2*. Here, we show that a loss of function *rin* mutation heterozygously suppresses eye degeneration caused by (G_4_C_2_)_30_, (GR)_36,_ or (PR)_36_ (**Fig. 7**), suggesting that loss of G3BP/Rin suppresses neurodegeneration in fly models of c9ALS/FTD.

**Figure 7:**
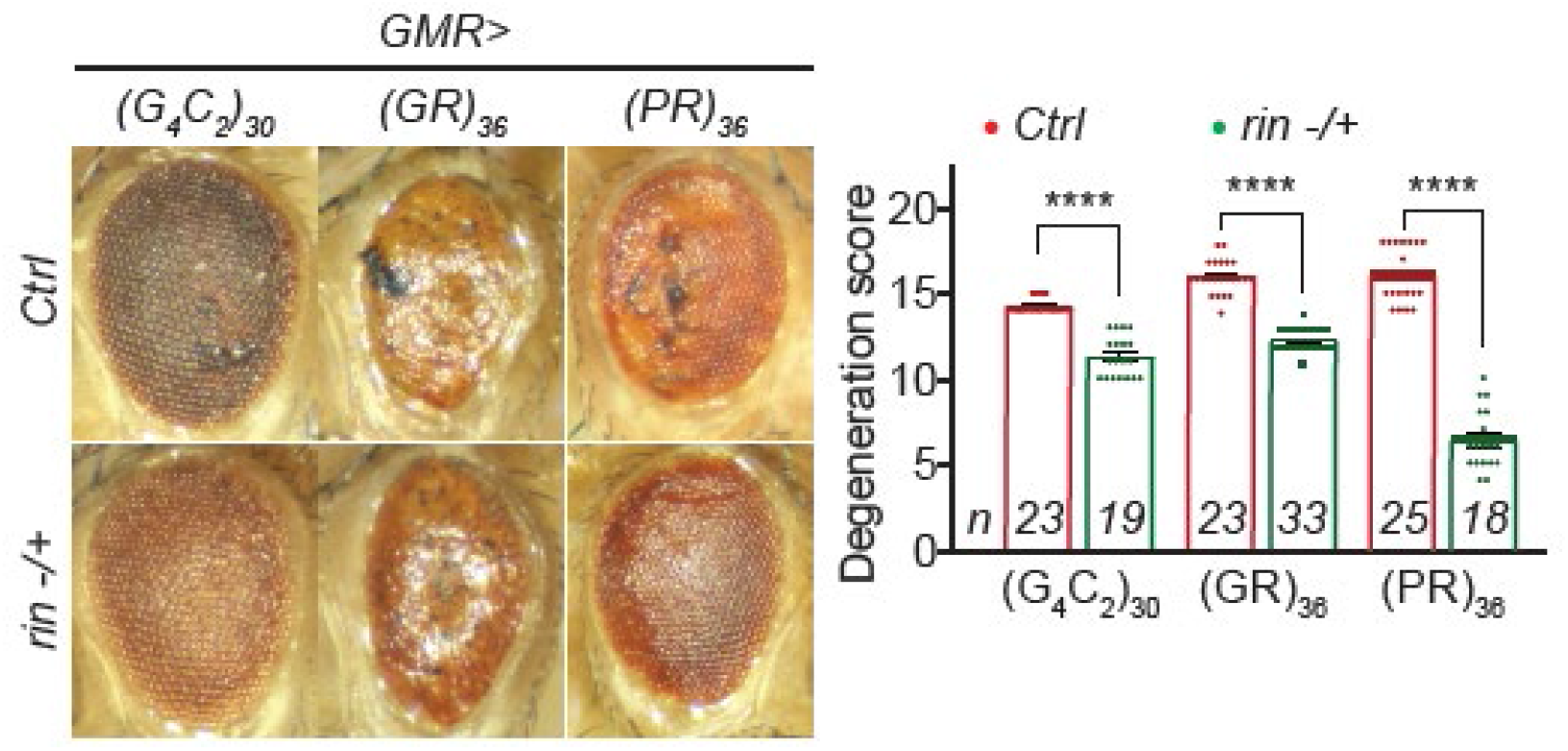
Loss of G3BP1/rin suppresses eye degeneration in c9ALS/FTD fly models. Fly eyes expressing (G_4_C_2_)_30_, (GR)_36,_ or (PR)_36_, without (Ctrl) or with heterozygous loss of function of *rin*. Mean ± s.e.m. Student’s t-tests; ****, *p*<0.0001.

## Discussion

Despite the importance of SGs in ALS/FTD pathogenesis, it is unclear how SG assembly is regulated at the cellular level and whether this regulation is related to pathomechanism. Here, we show that the ER stress/IRE1/JNK axis promotes SG formation caused by R-DPRs and contributes to neurodegeneration in animal and cellular models of c9ALS/FTD. Mechanistically, activated JNK promotes the *G3BP1* transcription, likely by phosphorylating H3S10, thereby increasing the G3BP1 protein level (**Fig. 8**). Together, our findings suggest a novel pathway regulating SG formation, which contributes to ALS/FTD pathogenesis.

**Figure 8:**
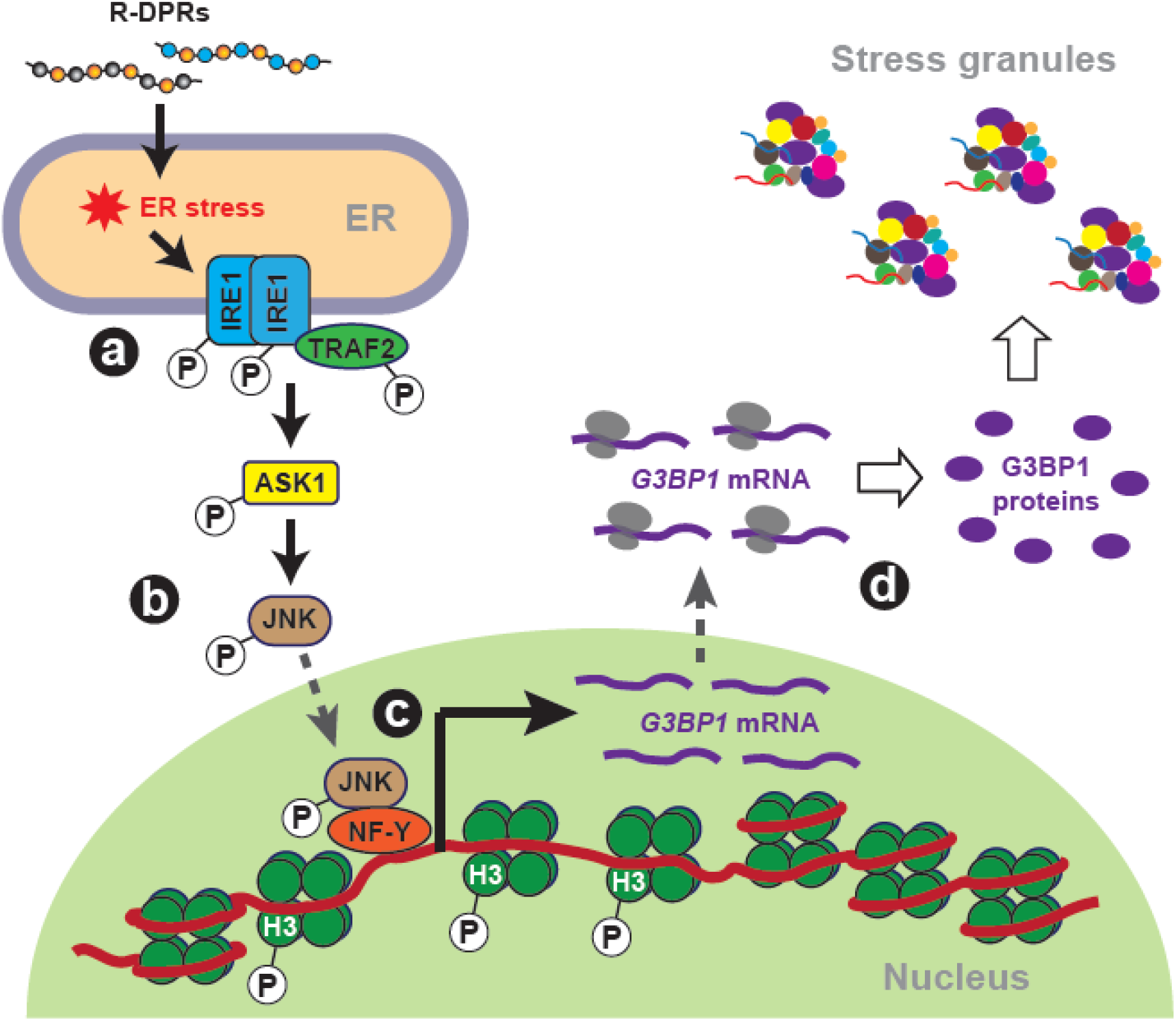
Schematic representation of ER stress/JNK promoting SG assembly in c9ALS/FTD. (a) R-DPRs induce ER stress, activating IRE1 and TRAF2. (b) Activated IRE1/TRAF2 complex activates ASK1, which subsequently activates JNK via phosphorylation. (c) Activated JNK translocates to the nucleus and, together with NF-Y, localizes to the promoter region of *G3BP1*, where it phosphorylates H3 at Serine10. H3S10 phosphorylation relaxes DNA, allowing NF-Y-mediated transactivation of *G3BP1*. (d) G3BP1 protein level is upregulated, causing SG assembly.

How JNK promotes G3BP1 transcription is unclear. A previous study showed that in mice neurons, the transcription factor complex nuclear factor Y (NF-Y) and active JNK are recruited to promoter regions of some genes, including *G3BP1* (Tiwari et al., 2011), where activated JNK phosphorylates H3S10, thereby allowing NF-Y-mediated *G3BP1* transactivation (**Fig. 8**). Future studies can test this model in U-2 OS and iPSN models of c9ALS/FTD.

When overexpressed, R-DPRs induce SGs in cells over a 24-48 hour period (Boeynaems et al., 2017; K. H. Lee et al., 2016) (and **Fig. 4** and **Supplementary Fig. 2**), whereas many stressors, e.g., arsenite, induces SGs within an hour. Previous studies on SG assembly mechanisms mostly focus on the latter, i.e., SGs induced by short-term stress (Jain et al., 2016; Kedersha et al., 2016; P. Yang et al., 2020). Some of the identified mechanisms are verified in SGs induced by long-term stress, e.g., eIF2α phosphorylation is required in both arsenite and R-DPR-induced SG formation (Boeynaems et al., 2017; K. Zhang et al., 2018). However, the uniqueness of long-term-stress-induced SG formation is unclear. Here, we show that JNK promotes SG formation induced by R-DPRs, but not one-hour treatment of arsenite, suggesting a mechanism specifically for long-term-stress-induced SG formation. This specificity likely comes from the ability of JNK to cause transcriptional changes, which, compared to posttranslational modifications, are more likely implicated in long-term stress responses (Q. Zhang et al., 2015).

Previous studies identified critical roles of the MAPK/JNK pathway in stress responses and neurodegeneration, including ALS. It induces apoptosis in a mouse model of SOD1-mediated ALS (S. Lee et al., 2016), causes energy deficiencies in a mouse model of Wallerian degeneration (J. Yang et al., 2015), and disrupts lipid metabolism due to mitochondrial oxidative stress in fly and mouse models (L. Liu et al., 2015). Our finding that JNK promotes SG formation in cellular models of c9ALS/FTD identifies a novel route by which this pathway contributes to stress responses and neurodegeneration, suggesting a broader role of MAPK/JNK.

In addition to the MAPK/JNK pathway, other pathways and processes are also known to contribute to SG formation, and targeting some of these pathways/processes suppresses neurodegeneration or cytotoxicity in ALS/FTD models (Becker et al., 2017; Gilks et al., 2004; Jain et al., 2016; Kedersha et al., 2016; Kedersha et al., 2008; Ohn et al., 2008; K. Zhang et al., 2018). However, the complex network regulating SG formation in cells is far from understood. Mass spectrometry analyses identified ∼400 proteins in yeast and mammalian SGs (Jain et al., 2016), and genetic screens identified more than 300 genes whose loss limits or reduces arsenite-induced SG formation in U-2 OS cells (Ohn et al., 2008; P. Yang et al., 2020). For most of these proteins/genes, how they contribute to SG formation and whether they are implicated in ALS/FTD pathogenesis is unclear. Future studies addressing these questions will provide a better understanding of SG biology and potentially identify novel therapeutic targets for the diseases.

## Materials and Methods

### IPSC culture and motor neuron differentiation

IPSC lines from C9orf72 patients and non-neurological controls were obtained from Cedars-Sinai Stem cell Core (patient demographics are provided in **Supplementary Table 1**). IPSCs were differentiated into direct induced motor neurons (diMNs) using a previously published protocol (A. N. Coyne et al., 2020). Briefly, iPSCs were grown in mTeSR media on Matrigel (Corning)-coated 10 cm dishes for two weeks before differentiation. At 40% confluency, iPSC colonies were cultured in Stage 1 media containing IMDM 47.5% (Gibco), 47.5% F12, 1% NEAA (Gibco), 1% Pen/Strep (Gibco), 2% B27 (Gibco), 1% N2 (Gibco), 0.2 μM LDN193189 (Stemgent), 10 μM SB431542 (StemCell Technologies), and 3 μM CHIR99021 (Sigma Aldrich) for six days. On Day 6, colonies were passaged with accutase (EMD Millipore) and re-plated on Matrigel-coated 6-well plates. Cells were cultured in Stage 2 media containing IMDM 47.5% (Gibco), 47.5% F12, 1% NEAA (Gibco), 1% Pen/Strep (Gibco), 2% B27 (Gibco), 1% N2 (Gibco), 0.2 μM LDN193189 (Stemgent), 10 μM SB431542 (StemCell Technologies), and 3 μM CHIR99021 (Sigma Aldrich), 0.1 μM all-trans RA (Sigma Aldrich), and 1 μM SAG (Cayman Chemicals) until Day 12. On day 12, cells were trypsinized (GenClone) and replated on Matrigel-coated 24-well plates for imaging or 6-well plates for biochemistry. Cells were cultured in Stage 3 media containing IMDM 47.5% (Gibco), 47.5% F12, 1% NEAA (Gibco), 1% Pen/Strep (Gibco), 2% B27 (Gibco), 1% N2 (Gibco), 0.1 μM Compound E (Millipore), 2.5 μM DAPT (Sigma Aldrich), 0.1 μM db-cAMP (Millipore), 0.5 μM all-trans RA (Sigma Aldrich), 0.1 μM SAG (Cayman Chemicals), 200 ng/mL Ascorbic Acid (Sigma Aldrich), 10 ng/mL BDNF (PeproTech), 10 ng/mL GDNF (PeproTech) until day 32. All cells were maintained at 37°C and 5% CO_2_.

### Propidium iodide (PI) staining

Day 32 diMNs were treated with 1 µg/mL of PI (Invitrogen) and one drop of NucBlue (Invitrogen) along with the media and incubated at 37°C and 5% CO_2_ for 30 min. Images were acquired using a Zeiss LSM 900 confocal microscope (Carl Zeiss) with an Axiocam 512 color camera and related software. For each condition, 10-15 images were taken.

### *Drosophila* genetics

*Drosophila* were raised on yeast-cornmeal-molasses food at 25°C. All RNAi fly stocks were procured from Bloomington Drosophila Stock Centre.

For eye degeneration assay, *GMR-Gal4, UAS-(G*_*4*_*C*_*2*_)_*30*_*/CyO* were crossed to Canton-S flies or *UAS-RNAi*, and *GMR-Gal4, UAS-(G*_*4*_*C*_*2*_)_*30*_*/+; UAS-RNAi (II or III)* and *GMR-Gal4, UAS-30R/+* were selected and aged at 27°C for 12 days. The external morphology of degenerated eyes was scored using a previously published method (G. P. Ritson et al., 2010). Briefly, points were added for necrotic patches, loss of bristles, retinal collapse, loss of ommatidial structure, and depigmentation of the eye. Both the eyes were scored and the individual scores were combined to give a total ‘degeneration score’ in the range of 0-20. Eye images were captured using a ZEISS SteREO Discovery.V8 microscope (Carl Zeiss) with Axiocam 512 color camera and related software.

For the lethality assay, male flies from *OK371-Gal4*; *UAS-(G*_*4*_*C*_*2*_)_*30*_*/Gal80, TM6* were crossed to virgin female Canton-S or *UAS-RNAi* flies at 25°C, and parent flies were removed after three days. After 15 days, non-tubby offspring, i.e. *OK371-Gal4/+; UAS-(G*_*4*_*C*_*2*_)_*30*_*/+; UAS-RNAi (II/III)/+*, was scored as either fully eclosed (live) or pharate lethal (unable to eclose).

For *bsk/JNK* activity in flies, male flies from *OK371-Gal4 or OK371-Gal4; UAS-(G*_*4*_*C*_*2*_)_*30*_*/Gal80, TM6* were crossed to virgin female flies from *puc-LacZ/TM6*. Third instar larvae of the offspring *OK371-Gal4/+; +/puc-LacZ or OK371-Gal4/+; UAS-(G*_*4*_*C*_*2*_)_*30*_*/puc-LacZ* were collected and their ventral nerve cords were dissected and subsequently stained.

For Xbp1-EGFP expression in flies, male flies from *OK371-Gal4* or *OK371-Gal4*; *UAS- (G*_*4*_*C*_*2*_)_*30*_*/Gal80, TM6* were crossed to virgin female flies from *Xbp1-EGFP*. Third instar larvae of the offspring *OK371-Gal4/+; +/Xbp1-EGFP* or *OK371-Gal4/+; UAS-(G*_*4*_*C*_*2*_)_*30*_*/Xbp1-EGFP*) were collected and VNC was dissected and stained.

### Cell culture

U2-OS cells (ATCC, HTB-96) were grown in DMEM (Gibco) supplemented with 10% fetal bovine serum (Gibco) and 1% penicillin-streptomycin and maintained at 37°C in a humidified incubator supplemented with 5% CO_2_. Transfections were performed using Lipofectamine 3000 (Invitrogen) reagent as per the manufacturer’s protocol. 48 h post-transfection, cells were either fixed and immunostained or lysed for immunoblot.

### Immunofluorescent staining

Fly VNCs were fixed with 3.7% formaldehyde for 20 minutes and penetrated in 0.4% PBX (PBS with 0.4% Triton X-100) for 1 hr at room temperature. Tissues were stained with primary antibodies anti-LacZ (DHSB, 40-1a, AB_528100,) and anti-elav (DHSB, 7E8A10) in 0.4% PBX and 10% donkey serum (DS) overnight. After that, VNCs were washed three times in 0.4% PBX (20 min each) and incubated with secondary antibodies conjugated to Alexa Fluor 488 and 568 (Thermo Scientific) in 0.4% PBX containing 10% DS. All primary antibodies were used at 1:200 dilutions and secondary antibodies were used at 1:1000 dilutions. Tissues were washed three times with 0.4% PBX (20 min each) and mounted on a coverslip using Prolong antifade Gold mountant (Invitrogen) along with DAPI.

U-2 OS cells or iPSNs were fixed with 4% paraformaldehyde for 20 min followed by penetration in 0.1% PBX (PBS with 0.1% Triton X-100) for 20 min at room temperature. For iPSNs, 0.3% PBX was used. Cells were blocked with 3% donkey serum (DS) followed by overnight incubation with primary antibodies in 0.1% TBST (TBS with 0.1% Tween-20) containing 3% DS. Primary antibodies were used as follows: G3BP1 (Abcam, 181149), G3BP2 (ProteinTech, 16276-1-AP, AB_2878237) and TIA1 (ProteinTech, 12133-2-AP, AB_2201427), at 1:200 dilutions. Cells were washed three times with TBST (20 min each) followed by incubation with secondary antibodies conjugated to either Alexa Fluor 488, 568, or 647 (1:1000 dilution) in TBST and 3% DS. After that, cells were washed thrice with TBST (20 min each) and mounted using Prolong antifade Gold mountant (Thermo Scientific).

Images were acquired using Zeiss LSM900 confocal microscope (Carl Zeiss) with an Axiocam 512 color camera and related software.

### Plasmid source and construction

The mCherry plasmid was procured from Addgene, and (GR)_100_-mCherry was a gift from Dr. Yong-Jie Zhang (Cook et al., 2020). To generate the mCherry-tagged poly-PR expression plasmid, the BioID sequence in myc-BioID-(PR)x100(F. L. Liu et al., 2022) was replaced with a NdeI/BamHI fragment encoding mCherry followed by a flexible linker (mCherry-GGGSx3).

### Drug treatments

U2-OS cells were treated with JNK inhibitor SP600125 (50 µM) (Selleck Chemicals) for 24 h or with IRE1 inhibitor 4µ8C (50 µM) (Selleck Chemicals) for 6 h and incubated at 37°C.

For iPSNs, day 32 diMNs were stressed with 5 µM of tunicamycin (Sigma Aldrich) and co-treated with either DMSO or 25-50 µM JNK inhibitor (SP600125) or 25-50 µM IRE1 inhibitor (4µ8C) for 24 h.

### Western blot, immunoblot

U2-OS and iPSNs were lysed in Laemmli buffer and heated to 98°C for 15 min. The protein samples were separated using 4-15% SDS mini-PROTEAN TGX precast gels (Bio-Rad) and transferred to 0.45 µm nitrocellulose membranes (Bio-Rad). For dot blots, samples were directly blotted on the nitrocellulose membrane and air-dried for 20 min. Blots were blocked with 5% milk for 1 hr and incubated overnight with the primary antibody (1:1000 dilution, for actin 1:5000) in 0.1% TBST containing 5% milk. Primary antibodies were used as follows: JNK (Cell Signaling, 9252S, AB_2250373,), phospho-JNK (Cell Signaling, 9251S, AB_331659), H3 (Cell signaling, 9715S, AB_331563,), phospho-H3S10 (Cell Signaling, 9706S, AB_331748), IRE1 (Cell Signaling, 3294S, AB_823545,), phospho-IRE1 (Abcam, ab48187, AB_873899), TRAF2 (Cell Signaling, 4724S, AB_2209845,), phospho-TRAF2 (Cell signaling, 13908S, AB_2798342), G3BP1 (Abcam, 181149), G3BP2 (ProteinTech, 16276-1-AP, AB_2878237) and TIA1 (ProteinTech, 12133-2-AP, AB_2201427) at 1:1000 dilutions and actin (EMD Millipore, MAB1501, AB_2223041,), mCherry (Abcam, ab167453, AB_2571870,) at 1:5000 dilutions. Blots were washed with TBST followed by incubation with HRP conjugated secondary antibodies in TBST and 5% milk (1:5000 dilution). For mCherry dot blots, bovine serum albumin (BSA) was used for blocking instead of milk. Chemiluminescent substrate WesternLigtning Plus-ECL (PerkinElmer) was used for detection. Images were captured using the iBright™ FL1500 Imaging system (Thermo Fischer Scientific).

### MTT assay

U2-OS cells were incubated with media containing MTT (1 mg/mL) at 37°C for 4 h. After that, media was removed and cells were lysed using DMSO, and absorbance was measured at 570 nm using A Tecan Spark multimode microplate reader.

### PCR

RNA was isolated from cells using Trizol reagent (Invitrogen) following the manufacturer’s protocol. Reverse transcription was done using the SuperScript IV First-Strand Synthesis kit (Invitrogen). Quantitative PCR was done using SYBR-Green Master mix using Applied Biosystem Quant Studio 7. Following primers were used: for *GAPDH* forward 5’-GTTCGACAGTCAGCCGCATC-3’, reverse 5’-GGAATTTGCCATGGGTGGA3-’; for *G3BP1* forward 5’-GTCCTTAGCAACAGGCCCAT-3’, reverse 5’-TTATCTCGTCGGTCGCCTTC-3’. To analyze *XBP1* alternative splicing events, cDNA was amplification using DreamTaq PCR mix (Invitrogen) and separated on 2.5% agarose gel (MetaPhor agarose, Lonza). Following primers were used: *XBP1* forward 5’-TTACGAGAGAAAACT CATGGC-3’, reverse 5’-GGGTCCAAG TTGTCCAGAATG C-3’.

### Quantification and Statistical analysis

Western blots were quantified using FIJI-just ImageJ. For SG counts and PI staining, at least 300 cells were counted. The data are presented as Mean ± SEM. Statistical analysis was done using paired or unpaired Student’s t-test, one-way ANOVA with Dunnett’s test, and Chi-square test using GraphPad Prism version 8 (GraphPad) as described in figure legends. * *p* < 0.05, ** *p* < 0.01, *** *p* < 0.001, *** *p* < 0.0001.

## Acknowledgments

We thank Yong-Jie Zhang and Karen Jansen-West for the constructs. Bloomington Drosophila Stock Center (NIH P40OD018537, fly stocks). Funding support: K.Z., NIH-NINDS/NIA (R01NS117461), DoD (W81XWH-21-1-0082), Target ALS, and the Frick Foundation for ALS.

## Supplementary figures

**Supplementary figure 1:**
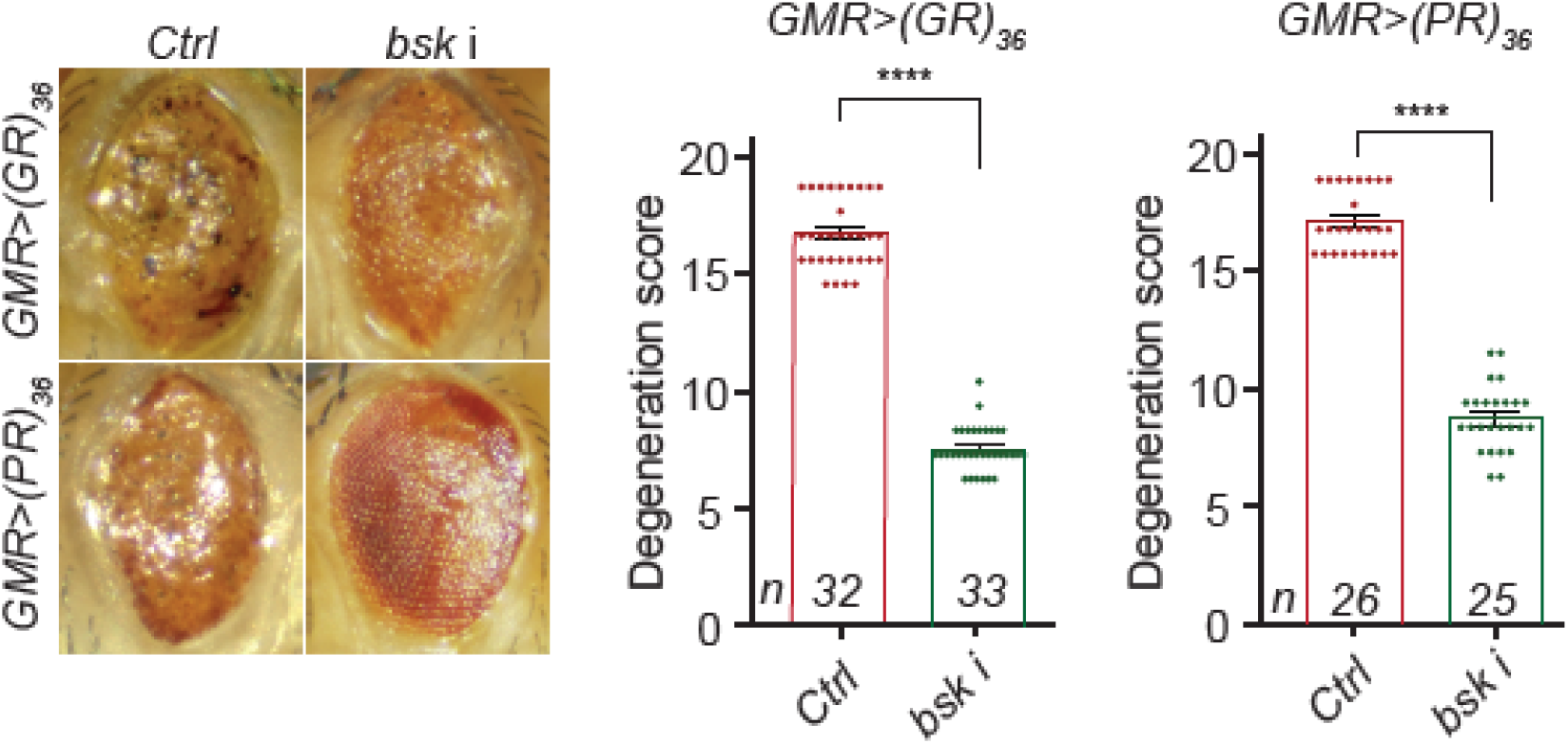
Loss of *bsk* suppresses R-DPR-mediated eye degeneration in flies. Fly eyes expressing (GR)_36_ or (PR)_36_ using GMR-GAL4, without (control, Ctrl) or with co-expressing *bsk* i. Student’s t-test; ****: *p*<0.0001

**Supplementary figure 2:**
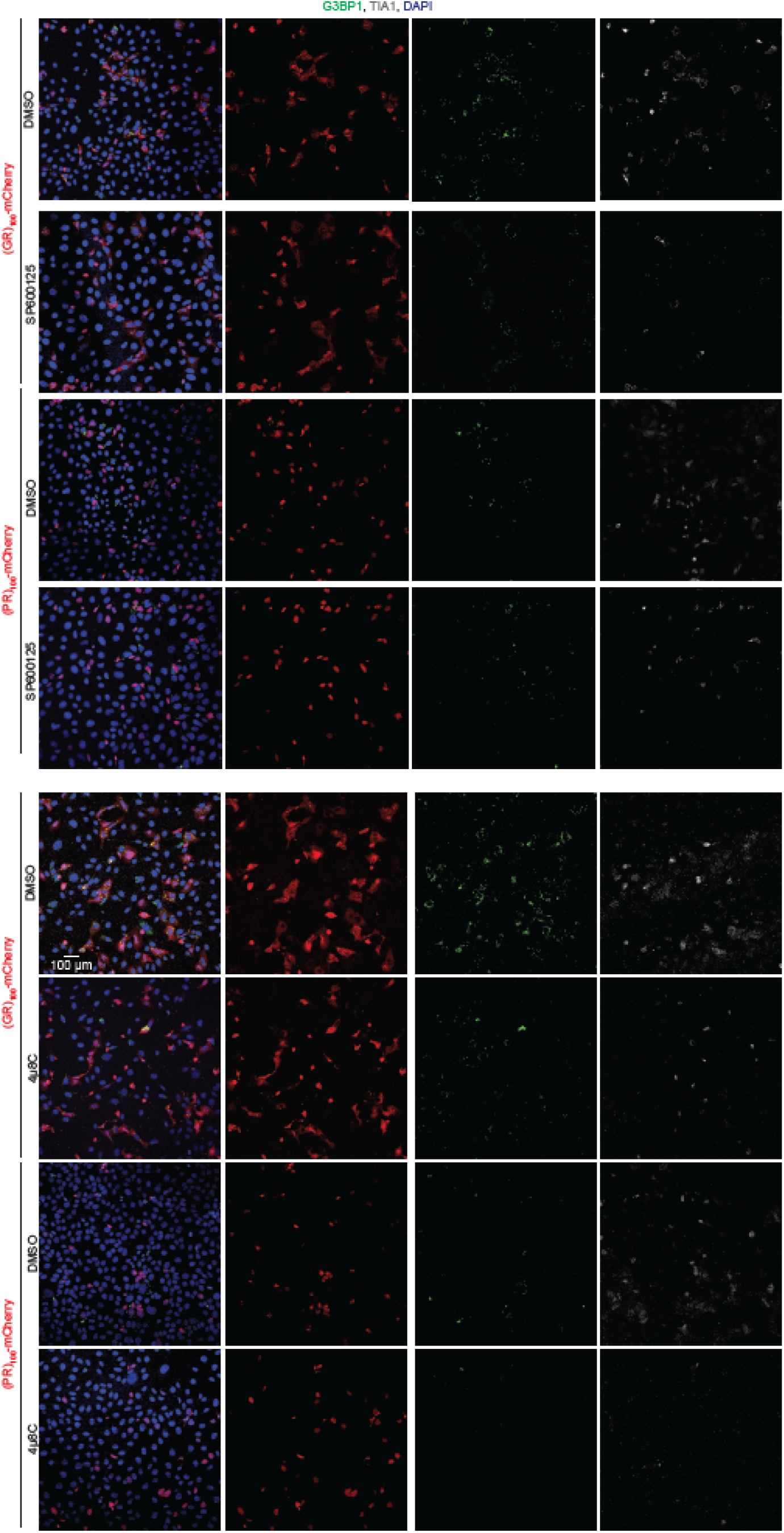
Large views of Fig. 4. U-2 OS cells expressing (GR/PR)_100_-mCherry (red) treated with DMSO, 50 µM SP600125, or 50 µM 4µ8C and stained with G3BP1 (green), TIA1 (white), and DAPI (blue).

**Supplementary figure 3:**
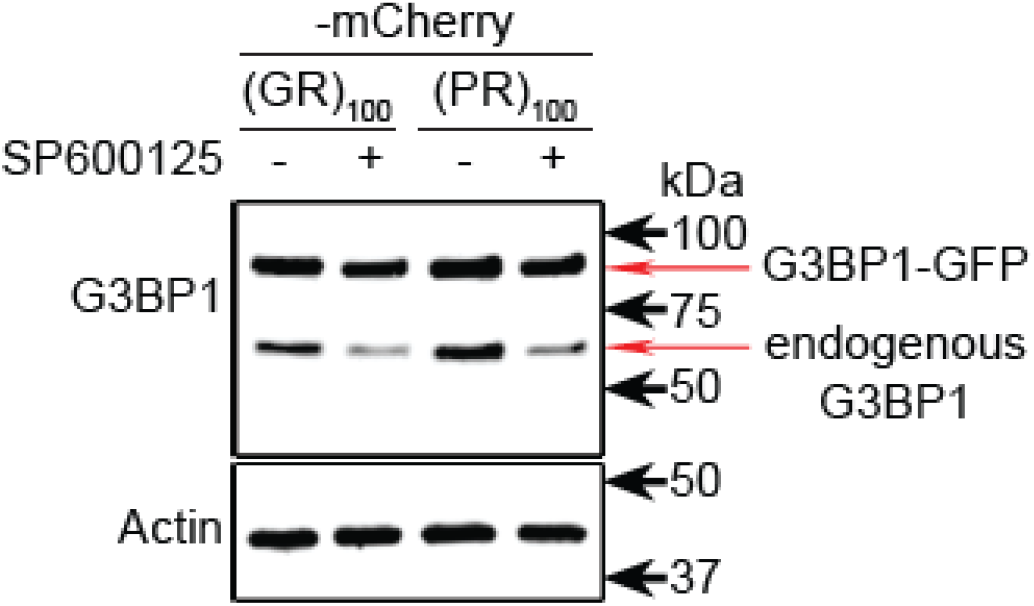
JNK regulates endogenous G3BP1. Western blots of lysates from U-2 OS cells stably expressing a G3BP1-GFP under a lentiviral promoter and transiently expressing (GR/PR)_100_-mCherry, treated with DMSO or 50 µM inhibitor SP600125.

**Supplementary figure 4:**
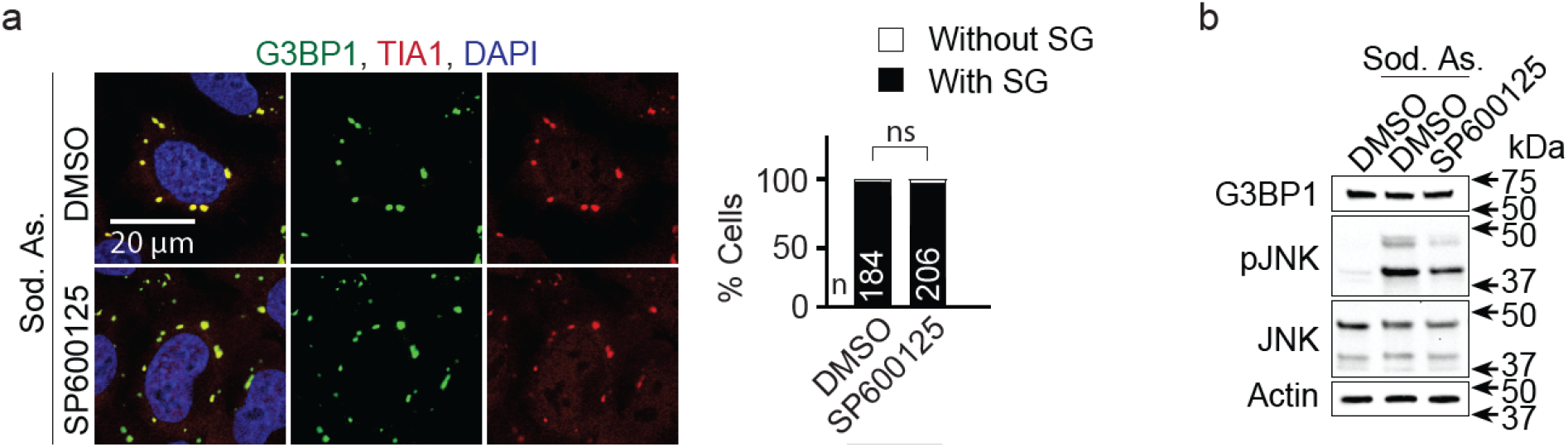
JNK does not promote arsenite-induced SG formation. (a) U-2 OS cells treated with 0.5 mM sodium arsenite (Sod. As.) together with DMSO or 50 µM SP600125 for 1 h and stained with G3BP1 (green), TIA1 (red), and DAPI (blue). Quantification shows percent of cells with or without SGs. Mean ± s.e.m. χ2-test. ns, not significant. (b) Western blots of lysates from U-2 OS cells treated with 0.5 mM sodium arsenite together with DMSO or 50 µM SP600125 for 1 h.

**Supplementary figure 5:**
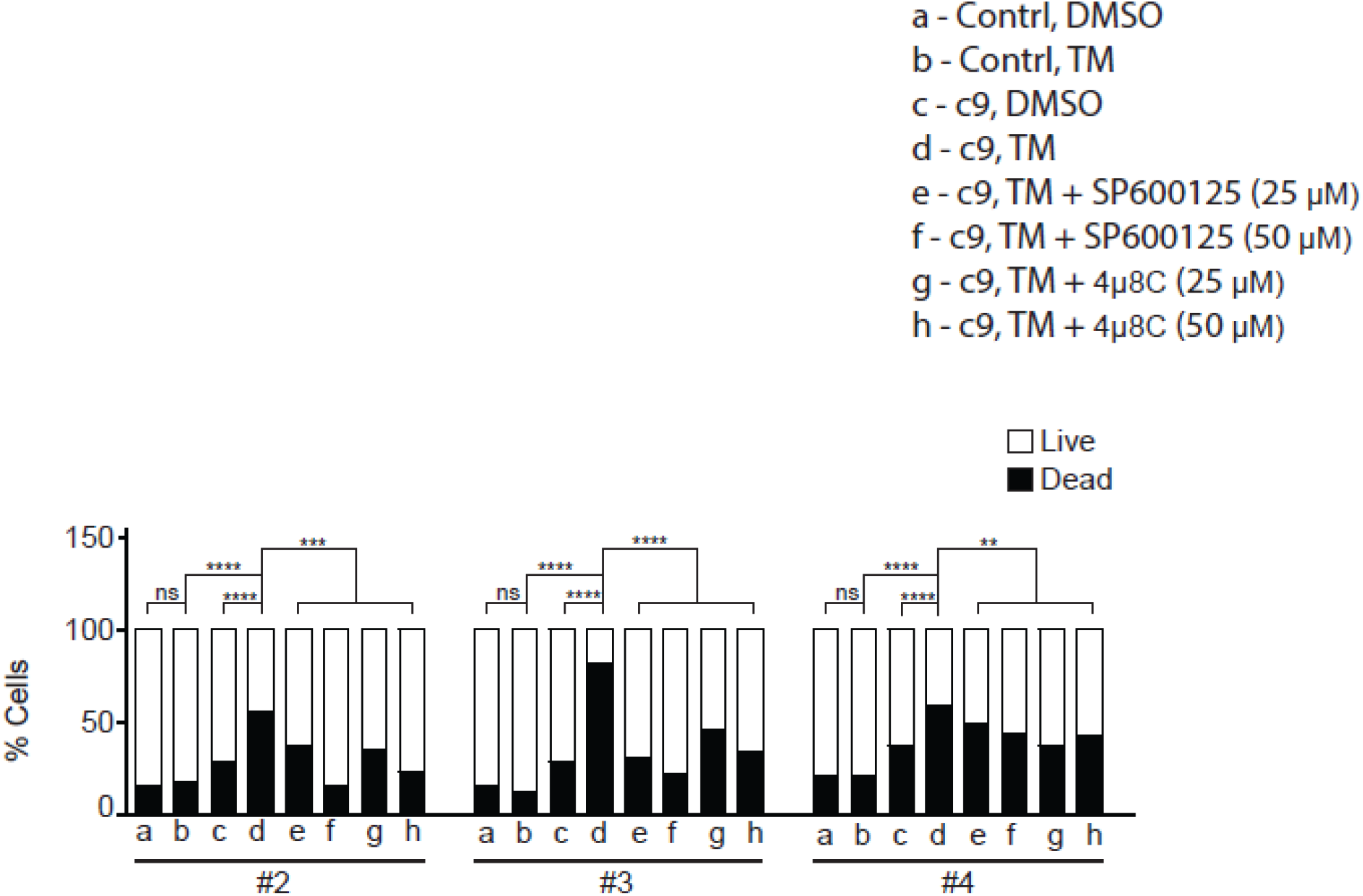
Inhibition of JNK or ER stress suppresses toxicity in multiple c9ALS/FTD iPSN lines. Control (Ctrl) or c9ALS/FTD (c9) Line #2, #3 and #4 iPSNs treated with 5 µM tunicamycin (TM) together with DMSO, SP600125, or 4µ8C and stained with propidine iodide (PI, dead cells) and NucBlue (all cells). Quantification shows percent of live and dead cells. χ2-test. ****: *p*<0.0001; ***: *p*<0.001; **: *p*<0.01; ns, not significant.

**Supplementary Table 1:**
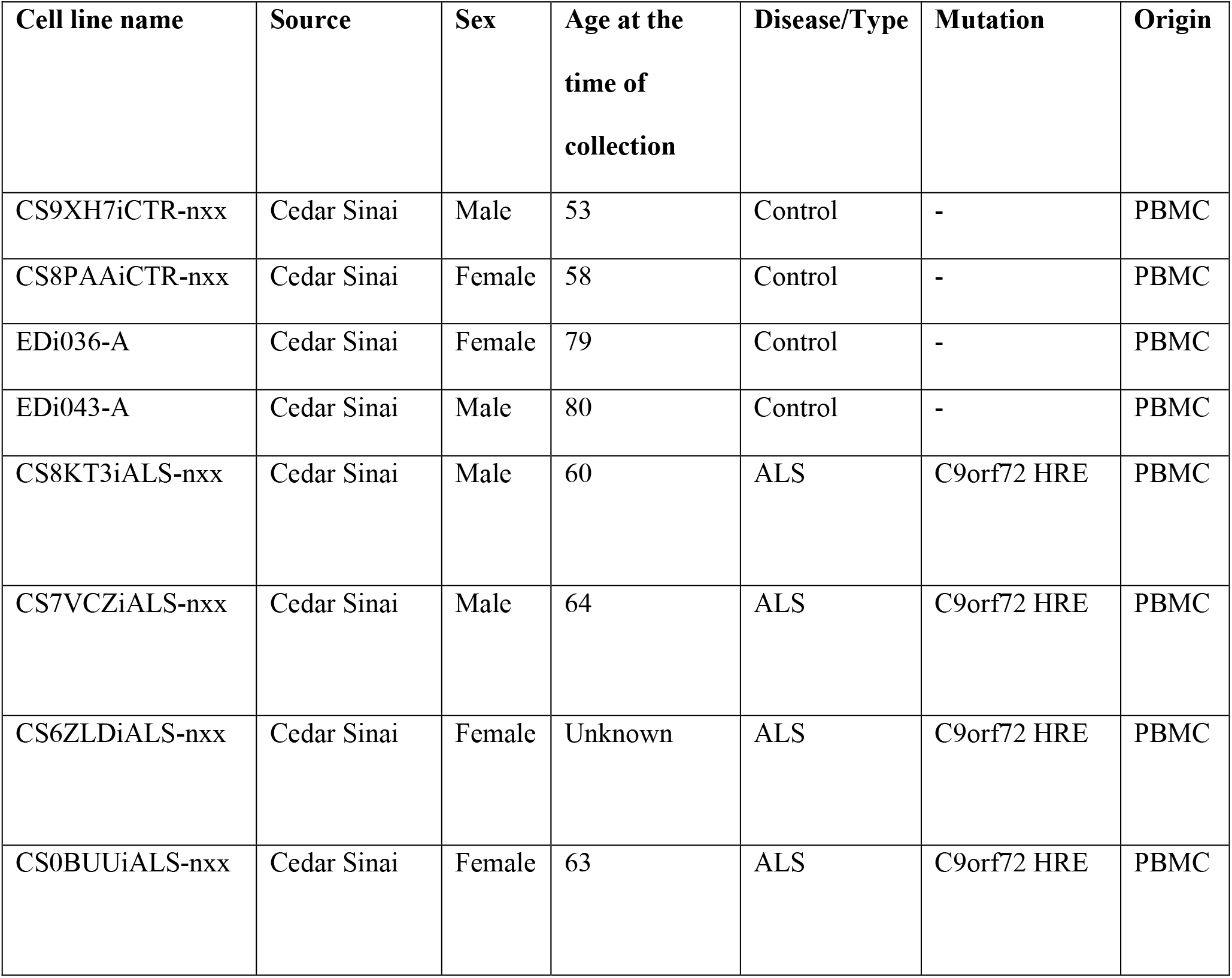
Demographics of patients whose iPSCs were used in this study.

| Cell line name | Source | Sex | Age at the<br>time of<br>collection | Disease/Type | Mutation | Origin |
| --- | --- | --- | --- | --- | --- | --- |
| CS9XH7iCTR-nxx | Cedar Sinai | Male | 53 | Control | - | PBMC |
| CS8PAAiCTR-nxx | Cedar Sinai | Female | 58 | Control | - | PBMC |
| EDi036-A | Cedar Sinai | Female | 79 | Control | - | PBMC |
| EDi043-A | Cedar Sinai | Male | 80 | Control | - | PBMC |
| CS8KT3iALS-nxx | Cedar Sinai | Male | 60 | ALS | C9orf72 HRE | PBMC |
| CS7VCZiALS-nxx | Cedar Sinai | Male | 64 | ALS | C9orf72 HRE | PBMC |
| CS6ZLDiALS-nxx | Cedar Sinai | Female | Unknown | ALS | C9orf72 HRE | PBMC |
| CS0BUUiALS-nxx | Cedar Sinai | Female | 63 | ALS | C9orf72 HRE | PBMC |

## Figure legends for source data

**Figure 1-source data 1:** Fly ventral nerve cord motor neurons expressing puc-lacZ without (G_4_C_2_)_30_, stained with lacZ (red), Elav (a neuronal marker, green), and DAPI (blue).

**Figure 1-source data 2:** Magnified inner image of fly ventral nerve cord motor neurons expressing puc-lacZ without (G_4_C_2_)_30_, stained with lacZ (red), Elav (a neuronal marker, green), and DAPI (blue).

**Figure 1-source data 3:** Fly ventral nerve cord motor neurons expressing puc-lacZ with co-expressing (G_4_C_2_)_30_, stained with lacZ (red), Elav (a neuronal marker, green), and DAPI (blue).

**Figure 1-source data 4:** Magnified inner image of fly ventral nerve cord motor neurons expressing puc-lacZ with co-expressing (G_4_C_2_)_30_, stained with lacZ (red), Elav (a neuronal marker, green), and DAPI (blue).

**Figure 2-source data 1:** Fly VNC motor neurons expressing Xbp1-GFP without (G_4_C_2_)_30_, stained with Elav (a neuronal marker, red) and DAPI (blue).

**Figure 2-source data 2**: Fly VNC motor neurons expressing Xbp1-GFP with (G_4_C_2_)_30_, stained with Elav (a neuronal marker, red) and DAPI (blue).

**Figure 3-source data 1**: U-2 OS cells expressing mCherry or (GR/PR)_100_-mCherry (red) stained with pJNK (green) and DAPI (blue).

**Figure 3-source data 2**: Raw image of Western blots for lysates from U-2 OS cell expressing mCherry or (GR/PR)_100_-mCherry showing actin

**Figure 3-source data 3**: Raw image of Western blots for lysates from U-2 OS cell expressing mCherry or (GR/PR)_100_-mCherry showing IRE1

**Figure 3-source data 4**: Raw image of Western blots for lysates from U-2 OS cell expressing mCherry or (GR/PR)_100_-mCherry showing p-IRE1

**Figure 3-source data 5**: Raw image of Western blots for lysates from U-2 OS cell expressing mCherry or (GR/PR)_100_-mCherry showing TRAF2

**Figure 3-source data 6**: Raw image of Western blots for lysates from U-2 OS cell expressing mCherry or (GR/PR)_100_-mCherry showing p-TRAF2

**Figure 3-source data 7**: Raw image of Western blots for lysates from U-2 OS cell expressing mCherry or (GR/PR)_100_-mCherry showing JNK

**Figure 3-source data 8**: Raw image of Western blots for lysates from U-2 OS cell expressing mCherry or (GR/PR)_100_-mCherry showing pJNK

**Figure 3-source data 9**: Raw image of Western blots for lysates from U-2 OS cell expressing mCherry or (GR/PR)_100_-mCherry co-treated with DMSO or 4µ8C for 6 h showing actin

**Figure 3-source data 10**: Raw image of Western blots for lysates from U-2 OS cell expressing mCherry or (GR/PR)_100_-mCherry co-treated with DMSO or 4µ8C for 6 h showing JNK

**Figure 3-source data 11**: Raw image of Western blots for lysates from U-2 OS cell expressing mCherry or (GR/PR)_100_-mCherry co-treated with DMSO or 4µ8C for 6 h showing pJNK

**Figure 5-source data 1**: Raw image of Western blots of lysates from U-2 OS cells expressing (GR/PR)_100_-mCherry, treated with DMSO or 50 µM SP600125 showing actin

**Figure 5-source data 2**: Raw image of Western blots of lysates from U-2 OS cells expressing (GR/PR)_100_-mCherry, treated with DMSO or 50 µM SP600125 showing G3BP1

**Figure 5-source data 3**: Raw image of Western blots of lysates from U-2 OS cells expressing (GR/PR)_100_-mCherry, treated with DMSO or 50 µM SP600125 showing G3BP2

**Figure 5-source data 4**: Raw image of Western blots of lysates from U-2 OS cells expressing (GR/PR)_100_-mCherry, treated with DMSO or 50 µM SP600125 showing H3

**Figure 5-source data 5**: Raw image of Western blots of lysates from U-2 OS cells expressing (GR/PR)_100_-mCherry, treated with DMSO or 50 µM SP600125 showing p-H3S10

**Figure 5-source data 6**: Raw image of Western blots of lysates from U-2 OS cells expressing (GR/PR)_100_-mCherry, treated with DMSO or 50 µM SP600125 showing TIA1

**Figure 5-source data 7**: Raw image of Western blots of lysates from U-2 OS cells expressing (GR/PR)_100_-mCherry, treated with DMSO or 50 µM 4µ8C showing actin

**Figure 5-source data 8**: Raw image of Western blots of lysates from U-2 OS cells expressing (GR/PR)_100_-mCherry, treated with DMSO or 50 µM 4µ8C showing G3BP1

**Figure 5-source data 9**: Raw image of Western blots of lysates from U-2 OS cells expressing (GR/PR)_100_-mCherry, treated with DMSO or 50 µM 4µ8C showing G3BP2

**Figure 5-source data 10**: Raw image of Western blots of lysates from U-2 OS cells expressing (GR/PR)_100_-mCherry, treated with DMSO or 50 µM 4µ8C showing H3

**Figure 5-source data 11**: Raw image of Western blots of lysates from U-2 OS cells expressing (GR/PR)_100_-mCherry, treated with DMSO or 50 µM 4µ8C showing p-H3S10

**Figure 6-source data 1**: Raw image of Western blot of c9 lysates treated with 5 µM tunicamycin (TM) together with DMSO, JNK inhibitor SP600125, or IRE1 inhibitor 4µ8C showing actin

**Figure 6-source data 2**: Raw image of Western blot of c9 lysates treated with 5 µM tunicamycin (TM) together with DMSO, JNK inhibitor SP600125, or IRE1 inhibitor 4µ8C showing G3BP1

**Supplementary Figure 3-source data 1**: Raw image of Western blots of lysates from U-2 OS cells stably expressing a G3BP1-GFP under a lentiviral promoter and transiently expressing (GR/PR)_100_-mCherry, treated with DMSO or 50 µM inhibitor SP600125 showing G3BP1

**Supplementary Figure 4-source data 1**: U-2 OS cells treated with 0.5 mM sodium arsenite (Sod. As.) together with DMSO for 1 h and stained with G3BP1 (green), TIA1 (red), and DAPI (blue).

**Supplementary Figure 4-source data 2**: U-2 OS cells treated with 0.5 mM sodium arsenite (Sod. As.) together with 50 µM SP600125 for 1 h and stained with G3BP1 (green), TIA1 (red), and DAPI (blue).

**Supplementary Figure 4-source data 3**: Raw image for Western blots of lysates from U-2 OS cells treated with 0.5 mM sodium arsenite together with DMSO or 50 µM SP600125 for 1 h showing JNK

**Supplementary Figure 4-source data 4**: Western blots of lysates from U-2 OS cells treated with 0.5 mM sodium arsenite together with DMSO or 50 µM SP600125 for 1 h showing pJNK

**Supplementary Figure 4-source data 5**: Western blots of lysates from U-2 OS cells treated with 0.5 mM sodium arsenite together with DMSO or 50 µM SP600125 for 1 h showing G3BP1

